# Exceptional diversity and selection pressure on SARS-CoV and SARS-CoV-2 host receptor in bats compared to other mammals

**DOI:** 10.1101/2020.04.20.051656

**Authors:** Hannah K. Frank, David Enard, Scott D. Boyd

## Abstract

Pandemics originating from pathogen transmission between animals and humans highlight the broader need to understand how natural hosts have evolved in response to emerging human pathogens and which groups may be susceptible to infection. Here, we investigate angiotensin-converting enzyme 2 (ACE2), the host protein bound by SARS-CoV and SARS-CoV-2. We find that the ACE2 gene is under strong selection pressure in bats, the group in which the progenitors of SARS-CoV and SARS-CoV-2 are hypothesized to have evolved, particularly in residues that contact SARS-CoV and SARS-CoV-2. We detect positive selection in non-bat mammals in ACE2 but in a smaller proportion of branches than in bats, without enrichment of selection in residues that contact SARS-CoV or SARS-CoV-2. Additionally, we evaluate similarity between humans and other species in residues that contact SARS-CoV or SARS-CoV-2, revealing potential susceptible species but also highlighting the difficulties of predicting spillover events. This work increases our understanding of the relationship between mammals, particularly bats, and coronaviruses, and provides data that can be used in functional studies of how host proteins are bound by SARS-CoV and SARS-CoV-2 strains.

## Main

The recent coronavirus pandemic has highlighted the disastrous impacts of zoonotic spillovers and underscores the need to understand how pathogens and hosts evolve in response to one another. Evolutionary analyses of host proteins targeted by infections reveal the pressures that hosts have faced from pathogens and how they have evolved to resist disease, informing predictions about spread of infections and how to counter them. The virus, SARS-CoV-2, the causative virus of COVID-19, like its close relative SARS-CoV, is thought to have its progenitor origins in bats^1–3^. Bats have been suggested to be “special” reservoirs of emerging infectious viruses^4^ and of coronaviruses in particular^5^. However, often this species-rich, ecologically diverse clade is treated as a homogenous group, represented by one or two species, particularly when considering the interaction of SARS-CoV and SARS-CoV-2 with host proteins (but see Hou et al.^6^ and Demogines et al.^7^ which consider multiple bat species).

Examination of host proteins bound by potential zoonoses can be used not only to infer past and current evolutionary pressure but to inform the likelihood of cross-species transmission. One major barrier to cross-species transmission is the ability of the virus, adapted to one host protein, to bind another species’ protein^8,9^. Accordingly, many studies have examined the ACE2 sequence of suspected disease reservoirs to understand how different viral strains bind different species’ ACE2 and where zoonotic spillover may have originated^6,8,10–12^. These studies, especially ones involving functional assays or in-depth modeling of virus-host contacts, are usually limited in their comparisons to a small subset of domestic animals or suspected reservoirs, e.g. rhinolophid bats or civets – a suspected intermediate host for SARS-CoV^13^. Often similarity, or lack thereof, between humans and other species in key ACE2 residues are used to predict the species that may have transmitted viruses to humans but because studies only examine a small subset of the existing diversity, it is hard to determine whether the selected species are more or less similar to humans than a random sample of animals.

Here, we investigate how angiotensin converting enzyme 2 (ACE2), the host protein bound by SARS-CoV and SARS-CoV-2^3,14,15^, has evolved in bats compared to other mammals. We analyze sequences drawn from 90 bat species, including 55 sequences generated for this study, an eight-fold increase over prior studies, and 108 other mammal species. Finally, we use our dataset of ACE2 sequences to highlight the potential for transmission of COVID-19 from humans into wildlife and the difficulty of predicting intermediate and amplifying hosts in spillover based on receptor similarity alone.

## Results and Discussion

### Mammals, particularly bats, are diverse at ACE2 contact residues for SARS-CoV and SARS-CoV-2

We analyzed a total of 207 ACE2 sequences from 198 species (90 bat species; 108 non-bat species) representing 18 mammalian orders (Table S1). There are 24 amino acid sites on ACE2 that are important for stabilizing the binding of ACE2 with the receptor binding domain of SARS-CoV (22 sites; Table S2) and/or SARS-CoV-2 (21 sites; Table S2)^6,8,10,11,14–16^. Across these 24 sites, which we refer to by their position in the human ACE2, we found a minimum of 132 unique amino acid combinations in the 207 sequences; across a subset of 7 amino acids identified as virus-contacting residues or important for the maintenance of salt bridges by most studies^6,8,15,16^ we found a minimum of 82 unique amino acid sequences (Figure 1). In bats (N=96), we found a minimum of 64 unique amino acid sequences across the 24 amino acids and 49 across the 7 amino acids, while in other mammals (N=111), we found a minimum of 68 unique amino acid sequences across the 24 amino acids and 38 across the 7 amino acids. Within species for which we were able to observe multiple individuals, we observed differing levels of diversity in the 24 sites with *Bos indicus, Rousettus leschenaultii, Camelus dromedarius* and *Rhinolophus ferrumequinum* identical within species across individuals and sites; *Canis lupus* showed one amino acid difference across two individuals; in contrast all four individuals of *Rhinolophus sinicus* were different from one another.

**Figure 1:**
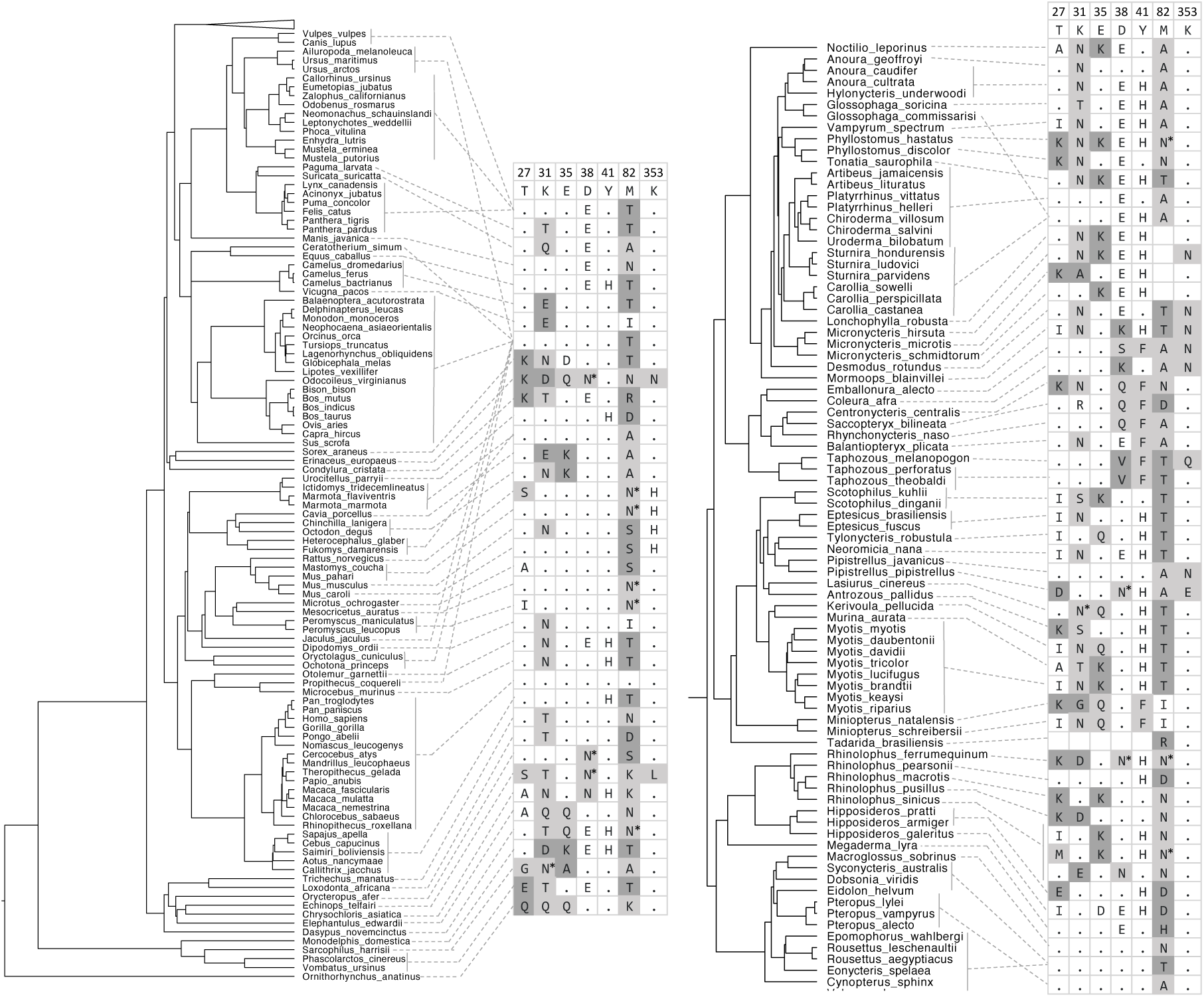
ACE2 is diverse across mammalian phylogeny at 7 residues responsible for contact with SARS-CoV and SARS-CoV-2. Dashed lines connect species to their ACE2 sequence. Numbers at the top of the table indicate the amino acid position in the human ACE2; human residues are listed as the reference. A period indicates identity with the human amino acid; unshaded boxes are amino acids that are identical or similar to humans; light gray indicates no or unknown impact on the interaction of ACE2 and SARS-CoV/ SARS-CoV-2 and dark gray indicates an amino acid that would likely disrupt the virus-host interaction. Asterisks on N indicate predicted presence of N-linked glycosylation. Blank boxes indicate a lack of data for that species and residue. The bat clade is collapsed on the left (top) and expanded in the right column.

Across all sequences, the 24 amino acid sites varied from monomorphic across all examined sequences (e.g. Phe^28^ and Arg^357^) to having 10 or more possible amino acids (e.g. 24, 27, 31, 34, 79, 82, 329). The most diverse sites (as measured by Shannon’s diversity index) were amino acid positions 24, 34, 79, 82, 329 and 354, while the most even sites were positions 24, 30, 34, 41, 82 and 329 (Table S2). Of the 22 sites with more than one amino acid, bats were more diverse than other mammals at 13 and were more even at 15. That bats demonstrate a similar diversity in their ACE2 across these loci and greater diversity in some sites than that observed across the rest of mammals suggests they may be particularly diverse in their ACE2, and supports the idea that bats are more diverse than other suspected SARS-CoV and SARS-like CoV hosts^6^.

### Bats drive the signal of mammalian selection and adaptation to SARS-CoV and SARS-CoV-2

We also conducted a series of selection analyses each on 5 phylogenetic trees drawn from Upham et al. ^17^. Across all mammals, the 20 variable sites in ACE2 that contact SARS-CoV were not more likely to be under positive selection than other residues in the gene (MEME p < 0.05, Fisher’s exact test, p_all trees_ >0.35; Table S3), though there was some marginal evidence for increased selection in these residues (MEME p < 0.1, Fisher’s exact test, p _two trees_ < 0.06; Table S3). Similarly, the 19 variable sites that contact SARS-CoV-2 were not more likely to be under positive selection than other residues in the gene when considering sites under selection at p < 0.05 (MEME p < 0.05, Fisher’s exact test, p_all trees_ > 0.1; Table S3). However, when considering sites under selection at p < 0.1, residues that contact SARS-CoV-2 do indeed appear to be more likely to be under selection than other residues in the gene, likely due to the reduction in statistical power loss (MEME p < 0.1, Fisher’s exact test, p_all trees_ < 0.02). Therefore there is some evidence that the locus is evolving in response to coronaviruses; this is similar to the finding of strong selection in aminopeptidase N (ANPEP) in response to coronaviruses in mammals^18^. However, this pattern is driven by and strengthened in bats; in bats a greater proportion of residues that contact SARS-CoV (MEME p<0.05, Fisher’s exact test, p_all trees_ < 0.03; MEME p<0.1, Fisher’s exact test, p_all trees_ < 0.02; Table S3) and SARS-CoV-2 (MEME p < 0.05, Fisher’s exact test, p_all trees_ < 0.02; MEME p<0.1, Fisher’s exact test, p_all trees_ < 0.0004; Table S3) were under selection than other residues in the gene, whereas residues that contact SARS-CoV (MEME p<0.05, Fisher’s exact test, p_all trees_ > 0.4; MEME p<0.1, Fisher’s exact test, p_all trees_ > 0.5; Table S3) or SARS-CoV-2 (MEME p<0.05, Fisher’s exact test, p_all trees_ > 0.4; MEME p<0.1, Fisher’s exact test, p_all trees_ > 0.5; Table S3) were not more likely to be under selection in non-bat mammals. Increased sampling can improve the ability of MEME to detect selection at individual sites^19^. Because our dataset of bat sequences is smaller than our mammalian dataset, it further strengthens our conclusion that bats are under positive selection in contact residues. Across all mammals, positions 24 and 42 were under selection (Table S2; 5 trees, MEME, p < 0.05), but in bats positions 27, 31, 35 and 354 (Table S2, 5 trees, MEME, p < 0.05) and 30, 38, 329 and 393 (Table S2, 5 trees, MEME, p < 0.1) were additionally under positive selection while positions 45 (Table S2, 5 trees, MEME, p < 0.05) and 353 (Table S2, 5 trees, MEME, p < 0.1) were under selection in non-bat mammals but not bats.

Using aBSREL we tested two *a priori* hypotheses, the first that bats are under positive selection in ACE2 and the second that the family Rhinolophidae, the bat family in which the progenitors of SARS-CoV and SARS-CoV-2 are hypothesized to have evolved^3,20^, specifically, is under positive selection. Both bats (p_all trees_ < 0.002) and Rhinolophidae (p_all trees_ < 0.00007) are under positive selection in ACE2 (Table S4). When we conducted an adaptive branch-site test of positive selection on all branches without specifying a foreground branch, branches in the bat clade were more likely to be selected than branches in other parts of the mammalian phylogeny (Fisher’s exact test, p_all trees_ < 0.05; Table S4) and the branch at the base of Rhinolophidae was under positive selection (p_all trees, Holm-Bonferroni correction_ < 0.04, Table S4). We found that bat branches are more likely to be under positive selection than other branches despite the fact that these branches are a subset of the total phylogeny and branch-site tests of positive selection have reduced power to detect selection on shorter branches, making our test conservative^18,21^. It is possible that the sequences we generated through target capture and genomic sequence were of poorer sequencing quality than the reference genomes (though the number of residues covered by sequences we generated and publicly available reference sequences was similar; two-tailed t-test, t = -0.49, p = 0.63). When we removed the bat sequences we generated and examined the remaining terminal branches, a greater proportion of bat branches were under selection than non-bat branches, but statistical significance was lost, likely due to reduced power (Table S4). Increased positive selection in bats in ACE2 compared with other mammals is consistent with their status as rich hosts of coronaviruses^5^. Host diversity of bats in a region is associated with higher richness of coronaviruses^5^; the diversity of bat ACE2 is consistent with the idea that a diversity of bats and their ACE2 sequences are coevolving with a diversity of viruses.

Two bat families, Rhinolophidae and Hipposideridae, have been associated with SARS-related betacoronaviruses^5^, which use the ACE2 molecule as a viral receptor^9^. Interestingly, while we found evidence that Rhinolophidae are under selection in ACE2, we found widespread selection across bats. Branches in the rhinolophid/ hipposiderid clade were not more likely to be under selection than other branches within bats (Fisher’s exact test, p_all trees_ > 0.7; Table S4) and bat lineages that live outside the predicted range of these viruses (e.g. in the Americas^5^) are also under positive selection. Therefore, there are still aspects of the bat-coronavirus relationship that we do not fully understand. At least one other coronavirus uses ACE2 to gain entry into the host cell, HCoV-NL63, which may have its origin in bats^22^; we found some evidence for increased selection in the residues that contact this virus in bats (MEME p < 0.05, Fisher’s exact test, p_all trees_ < 0.07; MEME p < 0.1, Fisher’s exact test, p_all trees_ < 0.08; Table S3), but not in non-bat mammals (MEME p < 0.05, Fisher’s exact test, p_all trees_ > 0.6; MEME p < 0.1, Fisher’s exact test, p_all trees_ > 0.4; Table S3). Many ACE2 residues that interact with HCoV-NL63 also interact with one or both of SARS-CoV and SARS-CoV-2^23,24^, which may be driving the evidence of selection in these residues. However, we did find evidence of selection in residues 321 and 326 in both bats and non-bat mammals (Table S2, 5 trees, MEME, p < 0.05), as well as selection in bats in residue 322 (Table S2, 5 trees, MEME, p < 0.05); these three residues contact HCoV-NL63 but not SARS-CoV or SARS-CoV-2. Our finding of selection in residues that contact HCoV-NL63 but not SARS-CoV or SARS-CoV-2 contrasts with the findings of a smaller dataset of bats mostly from Europe, Asia and Africa^7^ and may result from our greater power to detect signal or signal originating from bats in different regions than previously tested.

ACE2 regulation is known to impact survival in some influenza A infections^25^; New World bats are known to host many influenza A viruses^26^, so it is possible the selection we detect could result from infection from non-coronavirus infections. Still, our results raise questions about whether there are or were SARS-related coronaviruses in these regions that have not been detected?

### Similarity of ACE2 yields predictions of susceptible hosts but cannot completely determine host range of SARS-CoV and SARS-CoV-2

To determine how similar bats, civet, pangolin (a suspected source of SARS-CoV-2^27^) and other groups are to humans in 24 ACE2 residues that bind SARS-CoV (22 residues) and/or SARS-CoV-2 (21 residues), in all 207 sequences we quantified how many of the residues were identical or very similar to humans, likely maintaining current binding properties, versus how many were likely to disrupt binding. All of the apes and most of the Old World monkeys we examined were identical to humans across all amino acid sites; those that were not identical differed by only 1 or 2 amino acids (Figure 2; Table S1). However, amino acid similarity in these sites across different species often diverged from what we would have predicted from phylogeny alone. Notably, two rodents (*Mesocricetus auratus* and *Peromyscus leucopus*) had identical or very similar amino acids to humans in all but 2 sites for each virus, and many artiodactyls (e.g. cows, deer, sheep, goats), cetaceans, cats, and pangolin were as similar or more similar to humans than New World monkeys both in residues that contact SARS-CoV and in residues that contact SARS-CoV-2. The civet fell in the middle of mammals in its similarity to humans in residues that contact either or both viruses. In general, bats were not very similar to humans at these 24 amino acid sites, some with as many as five changes that would likely reduce virus binding, the most observed across mammals. Additionally, most bat sequences (56 of 91) showed that at least one of the two salt bridges (Lys^31^-Glu^35^; Asp^38^-Lys^353^ in humans) within ACE2 would be disrupted by changing a charged amino acid to an uncharged amino acid or to an amino acid with a clashing charge (Table S1). In Rhinolophidae, only one sequence of the ten examined did not have a change in position 31 or 35 that would result in a clash between two positively charged amino acids. Because of the large overlap in residues that contact SARS-CoV and SARS-CoV-2 (19 residues) generally species were roughly as similar to humans in residues that contact SARS-CoV and in residues that contact SARS-CoV-2 (Figure 2). However, bats (two-sided Wilcoxon signed rank test, p < 0.0001) and carnivores (two-sided Wilcoxon signed rank test, p < 0.0004), particularly mustelids including ferrets, were more similar to humans in residues that contact SARS-CoV-2 than residues that contact SARS-CoV (Figure 2).

**Figure 2:**
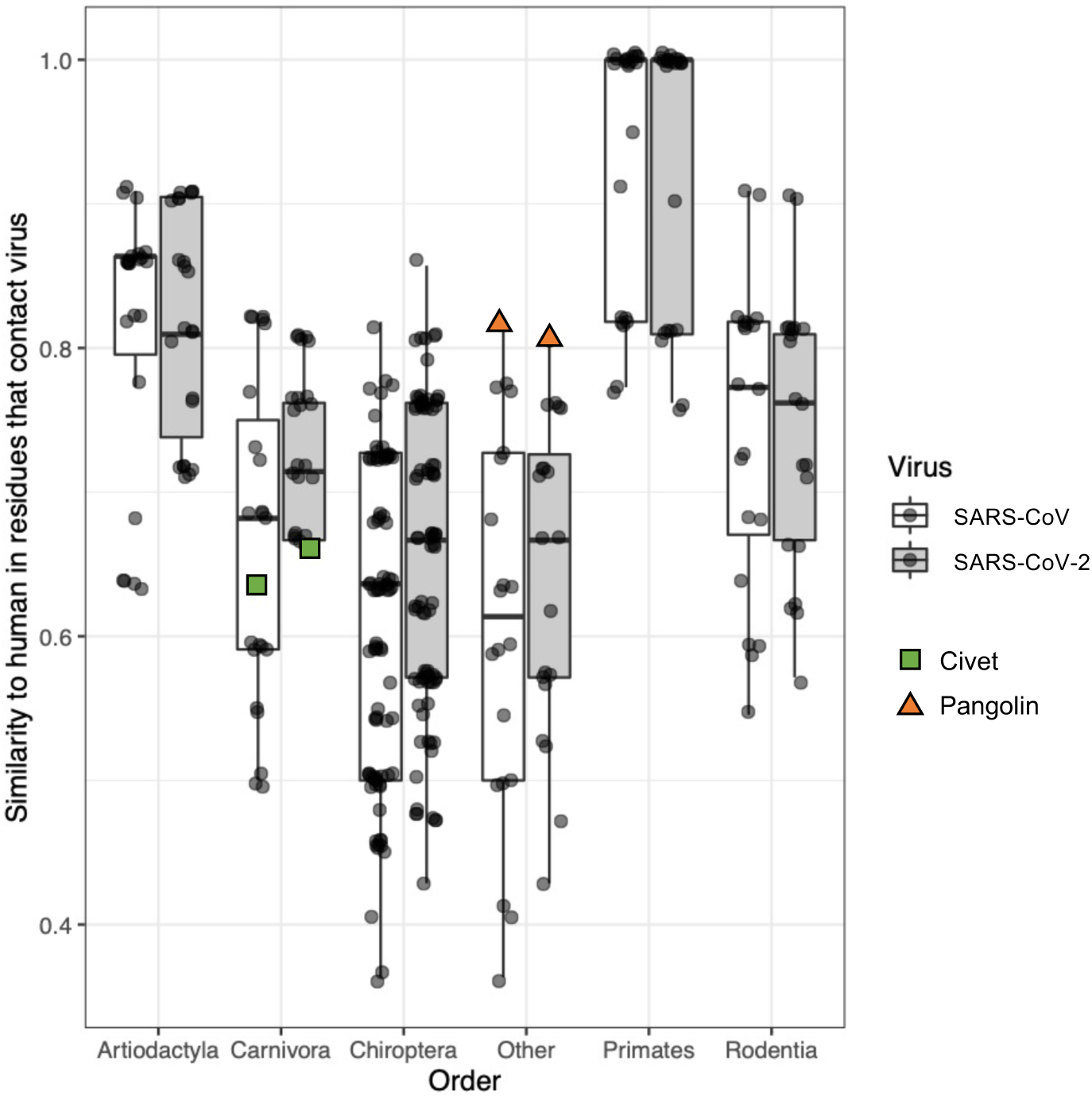
Similarity of ACE2 residues contacting SARS-CoV or SARS-CoV-2 to human. Similarity of residues was calculated based on the number of residues that were identical or highly similar in binding properties to those found in human ACE2 with penalties for residues that would likely disrupt binding (see methods). Scores of 1 indicate residues that contact the virus are identical (or highly similar to humans). Boxes cover the interquartile range with a line indicating the median and whiskers extending to the largest value less than 1.5 times the interquartile range. Each point indicates a single sequence; only sequences with data for at least 21 of 24 residues are shown. Sequences are grouped by mammalian order; “other” includes all orders with fewer than 4 sequences. The green squares in Carnivora indicate the civet (*Paguma larvata*) and the orange triangles at the top of the “Other” distribution indicate the pangolin (*Manis javanica*).

Examination of the diversity of ACE2 sequences across mammals and the similarity between distantly related groups at key residues for interaction with SARS-CoV and SARS-CoV-2 allows one to make predictions about potential spillover hosts or other susceptible species. In some cases, similarity of host residues seems to predict infection ability well. Old World primates were identical to humans across all 24 residues and, consistent with the idea that identical residues would confer susceptibility, experimental infections have demonstrated that SARS-CoV replicates in multiple macaque species^28^. Additionally, in our analysis, domestic cats were among the species most similar humans in residues that contact SARS-CoV and SARS-CoV-2. Notably, cats can become infected and can shed both SARS-CoV and SARS-CoV-2^29,30^. Pangolins were as similar in their ACE2 residues to humans as cats, lending some support for the idea that a virus that can bind pangolin ACE2 might be able to transmit to humans. Accordingly, it seems prudent to exercise precautions when interacting with species whose ACE2 is similar to humans in the contact residues for SARS-CoV and SARS-CoV-2, especially domestic animals such as cats, cows, goats and sheep. Care should also be taken with wild animals; for example, interactions between people with macaques or visitation of mountain gorillas by tourists could lead to cross-species transmission, endangering the health of humans and wildlife.

In other cases, it can be hard to predict susceptibility to SARS-CoV or SARS-CoV-2 infection based on overall similarity of ACE2 residues. A single amino acid change can impact the binding of a virus to ACE2. In position 24, a diverse, even and selected site across all mammals that contacts both viruses, mutation from Gln (human) to Lys (16 bat species and *Rattus norvegicus*) reduced association between the SARS-CoV spike protein and ACE2^12^.Position 27, a selected site in bats with many amino acid variants, is a Thr in humans; when mutated to a Lys (as in some bats), the interaction disfavored SARS-CoV binding by disrupting hydrophobic interactions with the SARS-CoV virus, while mutation to Ile, found in other bats, increased the ability of the virus to infect cells^6^. Some rodents, including *Mesocricetus auratus* and *Peromyscus leucopus*, which were among the most similar species to humans, have a glycosylated Asn in position 82 that disrupts the hydrophobic contact with Leu^472^ on SARS-CoV, reducing association between the spike protein and ACE2^12,15^; in conjunction with other mutations this glycosylation can disrupt binding^11^. We predict this same glycosylated Asn is also present in some rhinolophid bats (*R. ferrumequinum* and some *R. sinicus*), though not all (*R. pusillus, R. macrotis* and some *R. sinicus*). Additionally, intraspecies variation could be an important component of reservoir competency that we are unable to assess. It is likely that not all individuals in a species are equally susceptible to infection, complicating the identification of reservoirs.

Additionally, spillover potential is not regulated solely by ACE2 sequence and sometimes SARS-CoV or SARS-CoV-2 are able to replicate in hosts with divergent ACE2. Compatibility of the host protease with the virus is important for determining host range^9^ and viral strains vary in their binding properties to different species^15^, with some SARS-like coronaviruses able to bind human, civet and rhinolophid ACE2, despite major ACE2 sequence differences between the species^20^. Additionally, SARS-CoV can enter cells expressing the ACE2 of *Myotis daubentonii* and *Rhinolophus sinicus* with limited efficiency^6^, even though these species only share 13-16 amino acids with humans that contact either virus and each contain some mutations that should interfere with binding. Similarly, both SARS-CoV and SARS-CoV-2 replicated well in ferrets, whose ACE2 ranked among the least similar to humans in their contact residues for SARS-CoV, though more similar for SARS-CoV-2^29,30^. And species whose ACE2 sequence is not very similar to humans can be experimentally infected with SARS-CoV^11,31^.

## Conclusions

Taken together, our data suggest that mammals, particularly bats, are evolving in response to coronaviruses with a diverse suite of ACE2 sequences that likely confer differing susceptibility to various coronavirus strains. Predicting which species will expose humans to potential zoonoses is difficult. Data about viruses circulating in wildlife can help trace the origins of zoonotic disease outbreaks but are of minimal use if people continue to expose themselves to the reservoirs. Growing evidence suggests that some viruses are capable of evolving to infect different hosts, even when there might be sizable barriers such as divergent host receptors. The best solution is to prevent people and domestic animals from contacting wildlife to minimize opportunities for disease transmission and host switching.

## Acknowledgments

We thank Ellie Armstrong for assistance with bioinformatic analysis and Sandra Nielsen for insightful comments. We thank CONAGEBIO and the Organization for Tropical Studies for assistance and access to Costa Rican genetic resources. We thank the following museums for grants of tissue: Field Museum, Museum of Southwestern Biology, University of Alaska Museum, Museum of Vertebrate Zoology, University of Kansas Museum, and Texas Tech Museum. Additionally, we are grateful to the following organizations for funding the work: National Science Foundation Doctoral Dissertation Improvement Grant (1404521; HKF), Life Sciences Research Foundation Fellowship (HKF), Open Philanthropy Project, Stanford Woods Environmental Venture Program, Bing-Mooney Fellowship in Environmental Science, Stanford Center for Computational, Evolutionary and Human Genomics Postdoctoral Fellowship (HKF), National Institutes of Health grants “Molecular and Cellular Immunobiology” (5 T32 AI07290), Stanford School of Medicine Dean’s Postdoctoral Fellowship (HKF), and endowment from the Crown Family foundation (SDB).

## Author contributions

HKF, DE and SB planned the project. HKF collected and analyzed the data; DE contributed bioinformatic tools. HKF wrote the manuscript; all authors edited the manuscript.

## Data availability

The DNA sequences generated in this study are available from GenBank with the primary accession codes MT333480-MT333534.

## Code availability

No custom code was created for this analysis.

## Experimental methods

### Alignment of mammalian ACE2 sequences

Sequences for ACE2 were obtained either through Genbank or, in the case of several bat species, sequenced for this study. On 21 February 2020, ACE2 orthologs for all jawed vertebrates were downloaded from Genbank^32^. In addition, we sought out all bat sequences of ACE2, adding sequences from Hou et al.^6^, as well as the palm civet ACE2 sequences because of their putative role as reservoirs of SARS-CoV. Only sequences from mammalian species found in Upham et al.^17^ were retained and are listed in Table S1. Two Costa Rican bat species (*Artibeus* [*Dermanura*] *watsoni* and *Artibeus* [*Dermanura*] *phaeotis*) were not included in the phylogenetic hypotheses^17^ and were therefore excluded from the molecular evolution analyses but were included in analyses of amino acid identity and diversity. Multiple individuals were available for a few species (four individuals of *Rhinolophus sinicus*; three individuals of *Rhinolophus ferrumequinum*; and two each for: *Bos indicus, Camelus dromedarius, Canis lupus* and *Rousettus leschenaultii*), however only one sequence per species was used in molecular evolution analyses, noted in Table S1.

We sought additional data on the diversity of the ACE2 gene across bats using a combination of samples collected in the field in Costa Rica and granted from museums (63 species; summarized in table S1). For samples collected in the field, bats were captured in mist nets and a wing biopsy sample was collected. Bats were released immediately after sampling. Research was approved by the Stanford Institutional Animal Care and Use Committee (protocols 26920 and 29978) and conducted under the appropriate Costa Rican permits. From these species, we isolated DNA from tissue using the Qiagen DNeasy Blood and Tissue kit (Valencia, CA, USA) and created genomic libraries using the NEBNext Ultra II kit (New England BioLabs; Ipswich, MA, USA), according to manufacturer’s instructions. For some species, ACE2 was captured as part of a targeted capture using genomic libraries and a custom target enrichment kit (Arbor Biosciences; Ann Arbor, MI, USA) according to the manufacturer’s instructions with modifications^33^, while in other cases ACE2 was isolated from total genomic sequence. Briefly, genomic reads were mapped to the nearest bat genome of *Rousettus aegyptiacus, Pteropus alecto, Desmodus rotundus, Myotis lucifugus* or *Eptesicus fuscus* using LAST^34^ to generate a consensus sequence and the ACE2 coding regions were extracted using a translated DNA search in BLAT^35^ and the ACE2 coding sequence from *Myotis lucifugus*^36^. Sequences are available from Genbank (MT333480-MT333534; Table S1).

All sequences for ACE2 were aligned in Geneious^37^. Sequences were corrected by hand to remove sequences outside the coding region and adjust gaps to be in frame with the coding region using the human mRNA as a guide. Missing sequence, gaps and premature stop codons were converted to Ns for downstream analyses and comparison of residues with the human coding region.

### Investigation of important residues for CoV binding

We sought to understand how the residues important for coronavirus binding are conserved (or not) across mammals to determine probable host range of SARS-CoV and SARS-CoV-2. We compared amino acid sequences across 24 positions known to be important for binding of SARS-CoV and/or SARS-CoV-2 as determined by others^6,8,10,11,14–16^. To determine which amino acid positions were the most variable, we calculated Shannon’s diversity index (which accounts for the number and evenness of amino acids), number of unique amino acids and evenness for each of the 24 amino positions using the vegan^38^ package (version 2.5-6) in R^39^ (version 3.6.2). We also calculated how “human-like” a species was across these 24 amino acids, as well as separately for residues contacting SARS-CoV and residues contacting SARS-CoV-2 by giving a score to each amino acid in each position. Residues that were identical or relatively equivalent to humans were given a score of 1; relative equivalency was inferred when amino acids retained similar properties and abilities to participate in hydrogen bonds, Van der Waals forces or salt bridges. Residues that would likely be worse at binding were given scores of -1; reduced binding was inferred when amino acid properties were dramatically altered from that of the human amino acid motif (e.g. replacement of a positively charged amino acid with a negatively charged amino acid in a salt bridge). In general, asparagine and glutamine were considered similar enough not to disrupt binding, as were amino acids with the same charge and amino acids with small hydrophic side chains (valine, leucine, isoleucine and methionine). Amino acids whose effect was hard to determine were given scores of zero. Exact determinations of the impact can be found in Table S2. The human-like score was calculated as a sum of each amino acid score divided by the total amino acids observed across all 24 sites or all sites that contacted a given virus (since some species had missing data). We predicted the N-linked glycosylation of Asn when Asn was found in the following motif N-X-S/T where X is not a proline^40^. Glycosylation was not taken into account when calculating the human-like score.

### Molecular evolution analyses

To determine whether it was likely to be interactions with coronavirus driving the evolution of ACE2 we used MEME^19^ to infer the residues under selection across the mammal phylogeny, in just bats and in non-bat mammals and used a Fisher’s exact test to determine whether residues that interact with SARS-CoV, SARS-CoV-2 or HCoV-NL63^23^ were more likely to be under selection than other residues in ACE2. Only codons that showed variation (e.g. more than one amino acid across all 198 species) and that were present in humans were considered in the Fisher’s exact test. We used a p < 0.05 cutoff for inferring selection at each site via MEME but some results were shaper when using a p < 0.1 cutoff, likely due to the reduction in loss of statistical power (Table S3).

Additionally, to determine whether bats, and specifically the family Rhinolophidae, are under strong selection to adapt to viruses we used the adaptive branch-site random effects model test of positive selection, aBSREL^21^, as implemented in HyPhy, version 2.5.8^41^, using a pruned subset of five phylogenetic hypotheses chosen from the pseudo-posterior distribution of Upham et al.^17^ to account for phylogenetic uncertainty. We tested three conditions: 1) in which the branch leading to Rhinolophidae was considered the foreground branch; 2) in which the branch leading to the common ancestor of all bats was considered the foreground branch; and 3) in which all branches were tested without *a priori* specification of background and foreground branches. In determining whether bats are more likely to be under selection than other mammals, we used Fisher’s exact tests to test whether an excess of branches in the bat lineage were under selection compared to the rest of the phylogeny. We used p < 0.05 as our cutoff for a branch being under selection without any Holm-Bonferroni correction because it seemed unlikely that bat branches were more susceptible to false positives than any other branch and all our comparisons were between branches within the same aBSREL analysis. If one only accepts significance at a p < 0.05 with Holm-Bonferroni correction, a very stringent requirement, the general trends remain but the results lose statistical significance (Table S4). As described in the results, to guard against bias due to potentially lower sequence quality in the sequences we generated, we repeated our Fisher’s exact test using only terminal branches and removing sequences we generated; the trend of a larger proportion of bat branches being under selection was maintained but the results lose statistical significance (Table S4).

## Supplementary tables

Available as excel spreadsheets

**Table S1**: Data for each sequence on accession number, whether sequence is in selection analyses, preservation of salt bridges, identity of residues contacting SARS-CoV or SARS-CoV-2, combination of residues, and scores for similarity to humans in residues contacting SARS-CoV and SARS-CoV-2

**Table S2**: Summary data about residues that contact SARS-CoV, SARS-CoV-2 and HCoV-NL63 including diversity metrics, number of trees in which residues are inferred to be under selection, which virus is contacted by each residue and the identity of amino acids that lead to a positive or negative score in terms of similarity to humans.

**Table S3**: Results of Fisher’s exact tests on the number of selected and non-selected residues contacting SARS-CoV, SARS-CoV-2 and HCoV-NL63 as determined by MEME at p < 0.05 and p < 0.1. Results are summarized for tests on all 5 phylogenetic trees in all mammals, just bats or just non-bat mammals.

**Table S4**: Results of aBSREL analyses. Includes p-values from analyses in which the branch at the base of all bats is specified as the foreground branch, the branch at the base of Rhinolophidae is specified as the foreground branch or no foreground branches were specified. Also includes results of Fisher’s exact tests on the number of branches under selection in the bat clade compared to other branches; number of branches in the rhinolophid/ hipposiderid clade under selection compared to other branches in the bat clade; number of terminal bat branches (including or excluding sequences generated as part of the study) under selection compared to terminal branches in the rest of the phylogeny. Results using uncorrected p-values and Holm-Bonferroni corrected p-values are reported.

